# Evidence for an active handoff between cerebral hemispheres during target tracking

**DOI:** 10.1101/2025.04.18.649553

**Authors:** Matthew B. Broschard, Jefferson E. Roy, Scott L. Brincat, Meredith K. Mahnke, Earl K. Miller

## Abstract

The brain has somewhat separate cognitive resources for the left and right sides of our visual field. Despite this lateralization, we have a smooth and unified perception of our environment. This raises the question of how the cerebral hemispheres are coordinated to transfer information between them. We recorded neural activity in the lateral prefrontal cortex, bilaterally, as non-human primates covertly tracked a target that moved from one visual hemifield (i.e., from one hemisphere) to the other. Beta (15 to 30 Hz) power, gamma (30 to 80 Hz) power, and spiking information reflected sensory processing of the target. By contrast, alpha (10 to15 Hz) power, theta (4 to10 Hz) power, and spiking information seemed to reflect an active handoff of attention as target information was transferred between hemispheres. Specifically, alpha power and spiking information ramped up in anticipation of the hemifield cross. Theta power peaked after the cross, signaling its completion. Our results support an active hand-off of information between hemispheres. This handshaking operation may be critical for minimizing information loss, much like how mobile towers handshake when transferring calls between them.

## INTRODUCTION

The brain’s two hemispheres divide the world into left versus right visual hemifields. Each hemisphere has (somewhat) separate cognitive resources that are primarily devoted to the contralateral hemifield (Alvarez et al., 2012; Drew et al., 2008). This results in a “bilateral advantage,” such that attention is more successfully divided between multiple locations when they are in opposite hemifields than when they are within a single hemifield (Umemoto et al., 2010; Alvarez et al., 2005; Delvenne, 2005). Despite this lateralization, we perceive and act in a seamless fashion. This raises the question of how hemispheric resources are coordinated when items move from one side of vision to the other, as they often do.

Insights have come from tasks where targets are tracked as they move between the visual hemifields (Pylyshyn & Storm, 1988; Yantis, 1992; Weber et al., 2005; Bland et al., 2020; Drew et al., 2014; Landau et al., 2008; Chen et al., 2013; Holcombe et al., 2013; Holcombe et al., 2012). In human EEG, there seems to be an active handoff in which processing of a target overlaps in both hemispheres before and after it crosses the vertical meridian (Drew et al., 2014). The receiving hemisphere seems to anticipate the transfer, while the sending hemisphere continues to track the target after the transfer. This “handshaking” may reduce information loss caused by the transfer (e.g., transfer costs; Strong et al., 2020). When non-human primates (NHPs) are cued to transfer information held in visual working memory between hemispheres, there are increases in beta/gamma synchrony dynamics between the hemispheres, which is likewise suggestive of an active transfer process (Brincat et al, 2021).

We examined this process by recording neural activity from the lateral prefrontal cortex while NHPs covertly tracked visual targets that moved between hemifields. The transfer of targets was accompanied by oscillatory dynamics and spiking activity that seemed to anticipate and signal the transfer between hemispheres. Sensory events involved interplay between gamma (30–80 Hz), beta (15–30 Hz), and spiking information. Alpha (10–15 Hz) and spiking information anticipated the transfer. Theta (4–10 Hz) seemed to register the completion of the transfer.

## MATERIAL & METHODS

### Subjects

The non-human primate (NHP) subjects in our experiments were two adult male rhesus macaques (Macaca mulatta; 9 and 12 years old). Water was regulated to increase motivation. All procedures followed the guidelines according to the Massachusetts Institute of Technology Committee on Animal Care and the National Institutes of Health.

### Behavioral Paradigm

All procedures were controlled using custom-written MATLAB scripts and Monkey Logic software (Asaad et al., 2013). Eye movements were monitored using an EyeLink 1000 Plus (SR-Research; Ontario, Canada). The NHPs were positioned ∼50 cm from the computer screen (18” ASUS monitor; 1920 × 1080 resolution; 60k Hz refresh rate).

The NHPs maintained fixation on a white dot positioned at the center of the screen (Fig. 1A). After 300 ms of uninterrupted fixation, two gray dots were presented at 3.0 degrees eccentricity in one of two configurations: either the upper-left and lower-right quadrants (i.e., 135 and 315 degrees) or the upper-right and lower-left quadrants (i.e., 45 and 225 degrees; Fig. B). After 200 ms, one of the dots changed color (i.e., from gray to either red, blue, or green; the color identity was not relevant for this task). The NHPs were trained to attend to and covertly track the dot that changed color (i.e., the target), while ignoring the other dot (i.e., the distractor). After 200 ms, the color of the target changed back to gray, and the dots began rotating along a circular trajectory in the clockwise or counter-clockwise direction.

**Figure 1.**
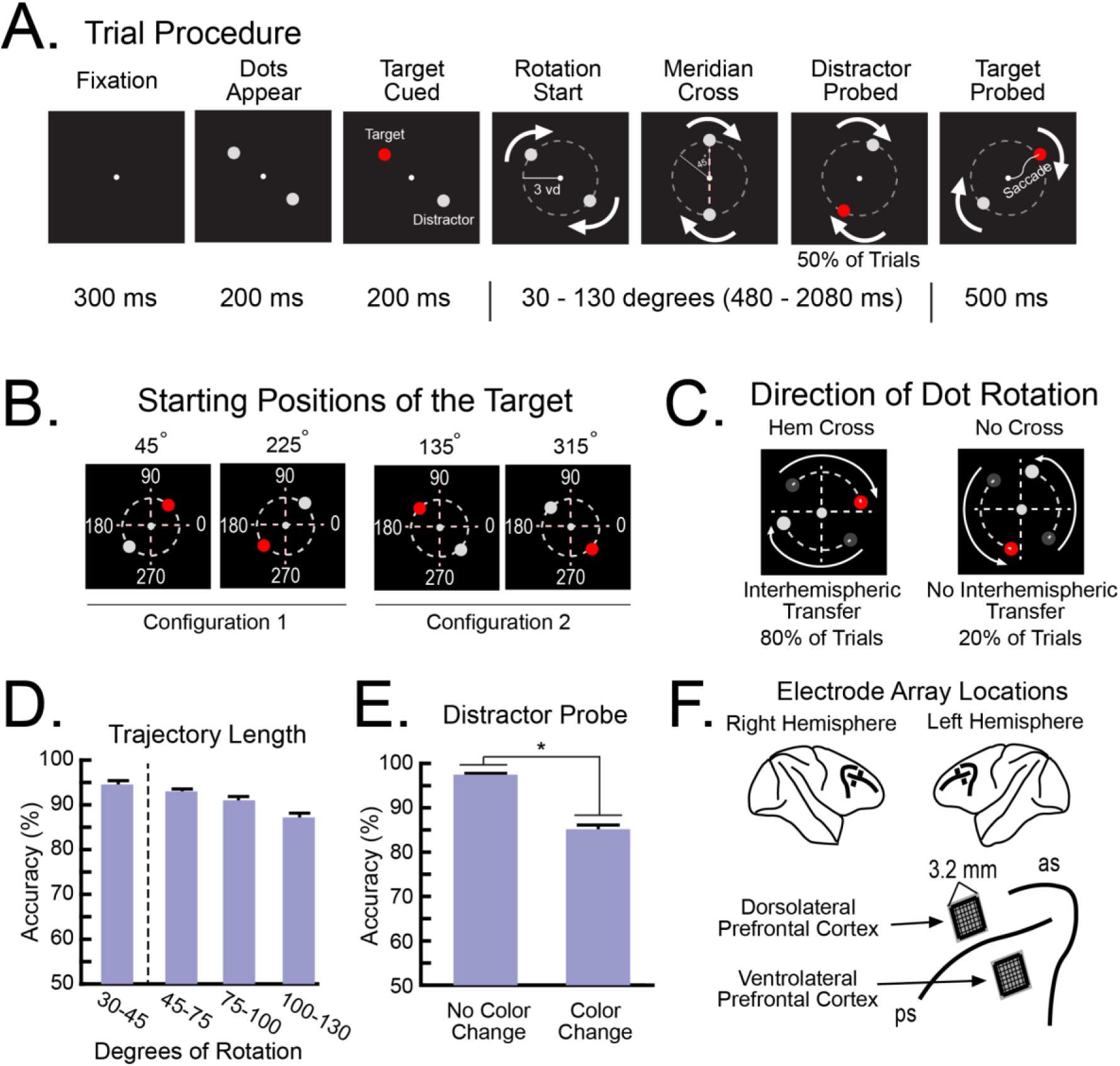
Methodology and behavior. **A,** Trial procedures. <u>Fixation:</u> The NHPs maintained fixation on a central fixation dot throughout the trial. <u>Dots Appear:</u> Two gray dots were presented on the screen. <u>Target</u> <u>Cued:</u> One of the dots changed color (e.g., gray to red). The NHPs were trained to attend to this dot (i.e., target) and ignore the other dot (i.e., distractor). <u>Rotation:</u> Both dots moved along a circular orbit in the clockwise or counter-clockwise direction. <u>Meridian Cross:</u> After 45 degrees of rotation, the dots crossed the vertical or the horizontal meridian. <u>Distractor Probed:</u> On 50% of trials, the distractor changed color at some point along the trajectory. The NHPs were trained to ignore this color change. <u>Target Probed:</u> After a random trajectory length, the target changed color again. The NHPs were trained to make a saccade to the target after it changed color. **B,** The dots appeared in two starting configurations (i.e., the two diagonals). Thus, the target started its rotation in one of the four corners (i.e., 45, 135, 225, or 315 degrees). **C,** For 80% of trials, the target crossed the vertical meridian (i.e., Hemispheric (Hem) Cross), requiring an interhemispheric transfer. For the remaining trials, the target crossed the horizontal meridian (i.e., No Cross) and did not require an interhemispheric transfer. **D,** Average performance during the Hem Cross trials. Accuracy decreased for trials with increasing trajectory length but remained high (>80%). The dotted vertical line separates trials with and without a meridian cross. **E,** Accuracy decreased for trials in which the distractor also changed color during the dots’ trajectory. **F,** Electrode arrays were implanted in the dorsolateral prefrontal cortex (dlPFC) and ventrolateral prefrontal cortex (vlPFC), bilaterally. ps = principal sulcus; ac = arcuate sulcus. All error bars indicate *S.E.M*. * indicates statistical significance.

Trials were categorized depending on whether the target crossed the vertical meridian (i.e., Hemispheric (Hem) Cross trials) or the horizontal meridian (i.e., No Cross trials; Fig. 1C). Both dots rotated at a constant rate of 16 ms per visual degree along the circular orbit. Therefore, the target crossed a meridian after 45 degrees of rotation (i.e., 720 ms). We biased the pseudorandom trial selection towards the Hem Cross trials (80% of trials), as these trials required an interhemispheric transfer, our main question of interest. The length of the dots’ trajectory ranged from 30 degrees to 130 degrees of rotation (480– 2,080 ms). The minimum trajectory length was shorter than the distance to the meridian. This ensured the NHPs were tracking the target before it crossed the meridian. The maximum trajectory length was chosen so that each trial never contained multiple hemifield crosses (i.e., a second cross would occur after 135 degrees of rotation).

At the end of the dots’ trajectory, the color of the target changed again (e.g., gray to red; Fig. 1A). The NHPs were given 500 ms to make a saccade from central fixation to the target. Successful saccades resulted in a juice reward. The distractor also changed color at a random location along its trajectory during 50% of the trials. The NHPs were trained to ignore this color change. This ensured the NHPs were attending to the target and ruled out the possibility that the NHPs were simply detecting *any* color change. Incorrect responses were defined as 1) a saccade to the distractor at any point along the trajectory or 2) a saccade to the target before it changed color. Trials were aborted if fixation was not maintained on the fixation dot for the duration of the trial.

### Electrophysiological Recordings

Four “Utah” microelectrode arrays (1.0 mm electrode length, 400 μm spacing; Blackrock Neurotech) were implanted into the lateral prefrontal cortex of each NHP, bilaterally. Arrays in the dorsolateral prefrontal cortex (dlPFC) were positioned ∼1 mm dorsal of the principal sulcus and ∼15 mm anterior to the genu of the arcuate sulcus. Arrays in the ventrolateral prefrontal cortex (vlPFC) were positioned ∼1 mm ventral to the principal sulcus and ∼10 mm anterior to the genu of the arcuate sulcus. Figure 1F shows the approximate positions of the arrays. Electrodes in each hemisphere were grounded and referenced to a separate subdural low-impedance reference wire. Local field potentials (LFPs) were amplified, low-pass filtered (250 Hz), and recorded at 30 kHz. The LFPs were then downsampled to 1000 Hz. Spiking activity was amplified, filtered (250–5,000 Hz), and manually thresholded to extract spike waveforms. Signals were recorded using the Grapevine interface (Ripple Neuro).

### Preprocessing

LFPs were re-referenced offline by subtracting off the mean signal from all electrodes in each array to remove common-source noise. Evoked potentials were removed from each electrode by subtracting off the mean signal, averaged separately for each of the target’s four starting locations. Outlier electrodes were defined as electrodes containing a grand average raw signal at least four standard deviations above the remaining electrodes within the array. These electrodes were removed from all analyses. Each analysis was calculated at each frequency between 4 Hz and 80 Hz and averaged into standard frequency bands (i.e., theta: 4–10 Hz; alpha: 10–15 Hz; beta: 15–30 Hz; gamma: 30–80 Hz).

Spiking activity from single neurons was sorted offline using the KiloSort 4.0 algorithm (Pachitariu et al., 2024) using the default parameters. Firing rates were calculated by summing the number of spikes into 50 ms non-overlapping bins. Neurons with a grand mean firing rate less than 1 Hz were excluded from all analyses.

Trials with a trajectory length less than 60 degrees were excluded from all analyses. This ensured that all analyzed trials had an adequate number of time points between the meridian cross and the color change of the target (i.e., at least 15 degrees of rotation, or 240 ms). Incorrect trials were also excluded from all analyses. Measures were normalized by dividing raw scores by activity 1.0 second before the initial fixation. We pooled trials with and without a color change of the distractor, as there were no differences in neural activity between these trial types (Supplemental Fig. 2).

Electrodes were defined as being either contralateral or ipsilateral to the target as it started its rotation (i.e., Sending and Receiving hemispheres, respectively). Within a given session, each electrode could be labeled as either a Sending or Receiving hemisphere, depending on the target’s starting location on each trial.

### LFP Analyses

#### Power

Power information was calculated by bandpass filtering the raw LFP signals (Butterworth filter) at each frequency. Then, the Hilbert transform was computed on the bandpass filtered data. Time-resolved power was calculated as the squared absolute value of this complex time series.

#### Coherence

LFP-LFP synchrony was estimated between electrode pairs by calculating coherence on the bandpass filtered LFP signal at each frequency. Coherence between two time series *X* and *Y* at frequency *f* was defined as

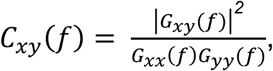

where *G_xy_* is the cross-spectral density between *X* and *Y*, and *G_xx_* and *G_yy_* are auto spectral densities of *X* and *Y*. All coherence values ranged from 0 to 1.

#### Granger Causality Analysis

Granger causality analysis was used to estimate the directionality of information between electrode pairs (Shojaie et al., 2022). Briefly, Granger causality uses regression to predict future values of a time series using past values. Time series X “Granger causes” time series Y, if adding past values of X improves the ability of the model to predict future values of time series Y, beyond the prediction based only on Y’s own past values. Granger causality was calculated using the *FieldTrip* MATLAB toolbox. The model order (number of past time points to include in the regression model) was fixed at 20 for all comparisons. This was determined by calculating Granger causality at different model orders (i.e., from 5 to 30 in intervals of 5) and comparing their model fits via Bayesian Information Criterion (BIC; Neath et al., 2011).

For this analysis, each time series was downsampled from 1000 Hz to 250 Hz to reduce the overall computational time. Additionally, we subsampled the number of electrode pairs. For each electrode in array ‘A’, Granger causality was calculated between that electrode and a randomly selected electrode in array ‘B’. Electrodes were selected without replacement to ensure signals from any given electrode were not sampled disproportionately.

#### Spike Decoding

Support vector machines (SVMs; Hearst et al., 1998) were used to assess whether the spiking activity carried information about the target’s location. Each classifier was trained on trials from one of the starting configurations (i.e., one of the two diagonals). For each trial, the classifier predicted which of the two dots on that diagonal was the target. The classifiers were trained with 80% of trials in each session and were tested with the remaining 20% of trials. Each classifier was trained using simultaneously recorded neurons in both hemispheres binned in 50 ms non-overlapping windows. To isolate the contribution of the Sending hemisphere and the Receiving hemisphere separately, we “ablated” the neurons from a single hemisphere (i.e. set all the corresponding classifier coefficients to 0) before measuring the classifier’s prediction on the testing set. We minimized bias in the model by averaging the classifier accuracies across fifty iterations. Different trials were used for the training and testing sets on each iteration. All models were fit using the *fitcsvm* function in MATLAB.

We used cross-temporal decoding (King et al., 2014) to examine how the neural codes generalized across time points. Identical procedures were used as above, except that each trained classifier was tested with spiking activity from all time points within the trial. This produces a two-dimensional matrix of classifier accuracies. Points along the diagonal reflect real-time decoding, where the same time point is used for the training and testing sets. Points off the diagonal reflect cross-temporal generalization, where different time points are used for the training and testing sets.

#### Statistical Analysis

For continuous neural measures that we could not assume were normally distributed, we used non-parametric statistics. Each neuron or LFP was treated as an independent observation. Randomizations were performed across all samples and were resampled

10,000 times to obtain a *p*-value. Repeated Measures ANOVA was used to assess changes in behavior.

## RESULTS

Two NHPs were trained to covertly track a target, one of two dots rotating around a computer screen (Fig. 1A). As the NHPs maintained a central gaze, a pair of dots appeared in opposite corners of the screen (Fig. 1B). The target was cued by one of the dots briefly changing color (Fig. 1A). The other dot was the non-target “distractor”. Then, the dots rotated along a circular orbit in either the clockwise or counter-clockwise direction. At a random time during this rotation, the target changed color again (Fig. 1A). The NHPs were rewarded for detecting this and making a direct saccade to the target. To ensure the NHPs were only attending to the target, the distractor also changed color along its trajectory on half of the trials (Fig. 1A). A saccade to it was an error.

As our study focused on how the target was transferred between hemispheres, most of the trials (80%) involved the target crossing from one hemifield to the other (right to left and vice-versa, “Hemispheric (Hem) Cross” trials; Fig. 1C). The hemispheres were characterized as being contralateral versus ipsilateral to the target at the start of the rotation (“Sending” hemisphere versus “Receiving” hemisphere, respectively). Our analyses focused on these trials. For a control comparison, on 20% of the trials, the target moved from the top to bottom or vice-versa within a single hemifield without a hemispheric transfer (“No Cross” trials; Fig. 1C).

The NHPs had high performance (∼90% correct; Fig. 1D) across all conditions (Supplemental Fig. 1A). Performance decreased for longer trajectory lengths (*F*(1,28) = 3.65, *p* = .026), but remained high for all lengths (>80% correct). Performance was about 10% lower on trials in which the distractor changed color during its trajectory (Fig. 1E; *F*(1,28) = 14.50, *p* < .001). During these error trials, we assume the NHPs were attending to the distractor. Accuracy was not significantly different between the Hem Cross trials and the No Cross trials (Supplemental Fig. 1B; *F*(2,27) = 2.96, *p* = .069). Spiking activity and local field potentials (LFPs) were recorded simultaneously from both hemispheres of the dorsolateral prefrontal cortex (dlPFC) and the ventrolateral prefrontal cortex (vlPFC) (Fig. 1F).

### Oscillatory dynamics reflect the transfer of the target between hemispheres

#### Higher frequencies in vlPFC reflected sensory events

Gamma and beta power, especially in the vlPFC, reflected sensory events. Figures 2A & B show heatmaps of average power in each hemisphere during the Hem Cross trials. In the vlPFC, gamma power peaked during the initial fixation and when the dots appeared (Fig. 2C right). These peaks occurred in both the Sending and Receiving hemispheres equally, since sensory stimulation was similar in both hemifields during these trial events. A third peak in gamma power occurred when the target was cued (Fig. 2C right). This peak was only observed in the Sending hemisphere, which was contralateral to this cueing event (Sending hemisphere versus Receiving hemisphere: *p* < .001). These gamma effects were similar in the control No Cross trials, which had similar sensory stimulation during these trial events (Fig. 2C right). Beta power in the vlPFC showed a similar pattern of effects to the sensory events, albeit in the negative direction. During both Hem Cross trials and No Cross trials, beta power tended to decrease as gamma power increased (Fig. 2D right). In fact, beta and gamma power were anti-correlated during these trial events (Supplemental Fig. 3). All effects were stronger in the vlPFC than the dlPFC (Figs. 2C & D left; all *p* < .01), consistent with previous work showing that the dlPFC is less activated by sensory inputs (Wutz et al., 2018). These results suggest that vlPFC gamma power is primarily driven by external, bottom-up sensory events.

**Figure 2.**
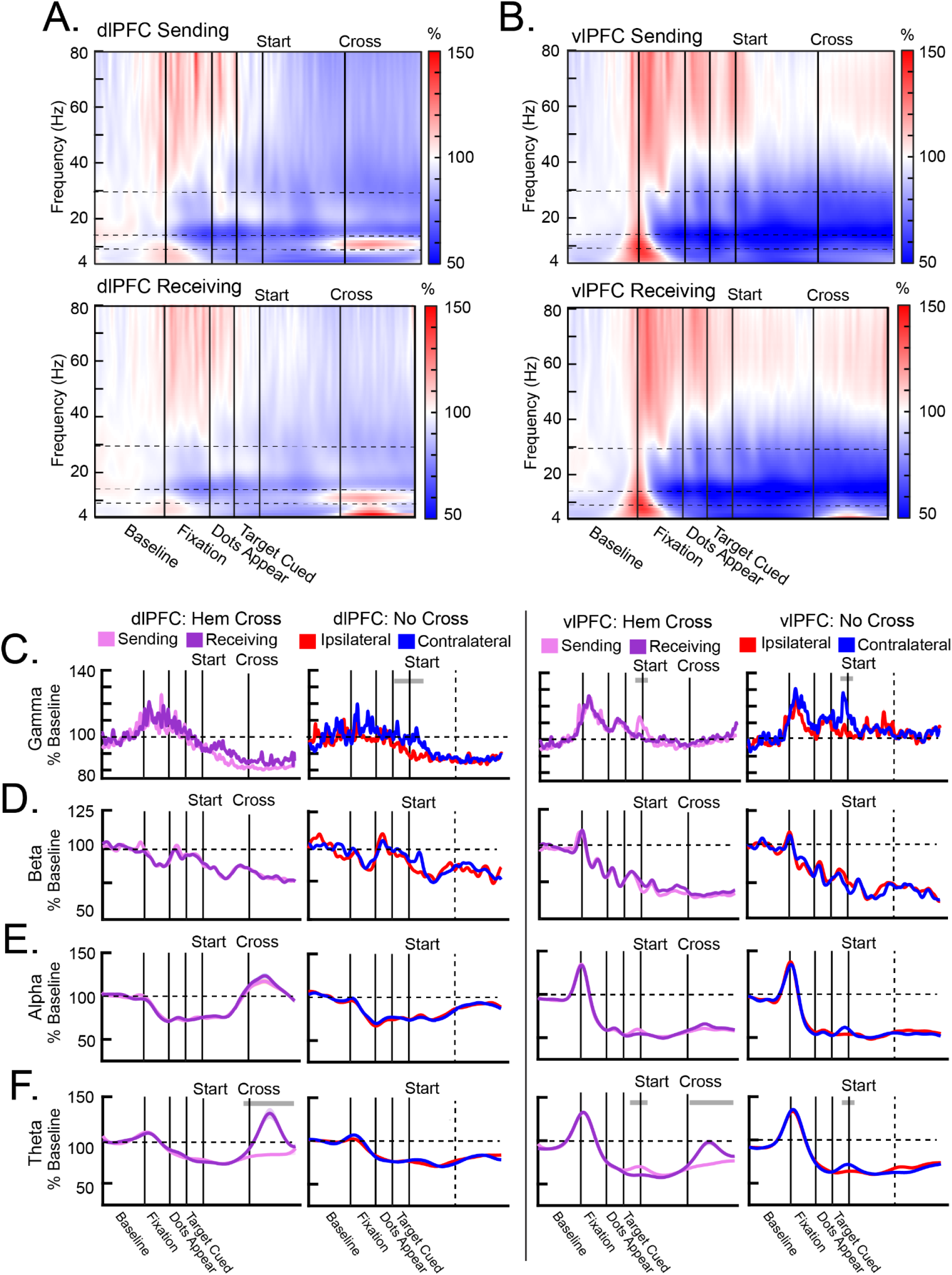
LFP power reflected sensory processing and transfer dynamics. **A–B**, Heatmaps of averaged power in the dlPFC (**A**) and vlPFC (**B**) during the Hemispheric (Hem) Cross trials, separated by the Sending hemisphere (top) and the Receiving hemisphere (bottom). Horizontal dotted lines indicate the different frequency bands. **C-F,** Power averaged across the gamma (**C**), beta (**D**), alpha (**E**), and theta (**F**) frequency bands in the dlPFC (left column) and the vlPFC (right column), separated by the Hem Cross trials and the No Cross trials. The vertical dotted lines in the No Cross plots indicate when the target crossed the horizontal meridian. All error bars indicate *S.E.M.* Gray bars above each plot indicate time points in which power was significantly different between hemispheres. Beta and gamma power in the vlPFC tracked the sensory events. dlPFC alpha power ramped up in anticipation of the cross. dlPFC theta power in the Receiving hemisphere peaked after the cross.

#### Lower frequencies in dlPFC reflected the interhemispheric transfer

Alpha power in the dlPFC increased in the lead up to the target’s cross from one hemifield to the other (Fig. 2E left). Starting about 250 ms before the cross, there was significant ramping of alpha power in both the Sending and Receiving hemispheres (Fig 2E left; slope of alpha power was greater than 0; all *p* < .001). This peaked shortly after the target crossed hemifields. By contrast, there was significantly less ramping during the No Cross trials when the target moved across the horizontal meridian (Fig. 2E left; difference in slope; *p* < .001), indicating that it was specifically related to information transfer between hemifields. This effect was only present in the dlPFC and was not evident in the vlPFC (Fig. 2E right).

Theta power, primarily in the dlPFC, peaked after the target crossed hemifields (Fig. 2F left). This peak only occurred in the Receiving hemisphere, which was contralateral to the target after the hemifield cross (Sending versus Receiving hemispheres: *p* < .001). This effect also did not occur in the No Cross trials, indicating it was also specific to the interhemispheric information transfer. Similar to alpha, the effects in theta were stronger in the dlPFC than the vlPFC (Fig. 2F right; all *p* < .05). Together, these results suggest that theta and alpha power in the dlPFC are involved in top-down control mechanisms that mediate the transfer of information between visual hemifields, and thus between cortical hemispheres.

### Interhemispheric connectivity was largely bidirectional

Granger causality (Fig. 3) and coherence (Supplemental Fig. 4) between hemispheres largely mirrored the changes in power. Figures 3A & B show heatmaps of Granger causality between the Sending and Receiving hemispheres (Fig. 3A), as well as the contrast between these directionalities (Fig. 3B). Granger causality indicated largely bi-directional influences between hemispheres. The exception to this occurred when the target was cued. At this time, influences in the vlPFC flowed from the Sending hemisphere (contralateral to the cue) to the Receiving hemisphere (Figs. 3C–F). This directionality was evident across all frequency bands but was especially prominent in beta and theta (Figs. 3B, D & F).

**Figure 3.**
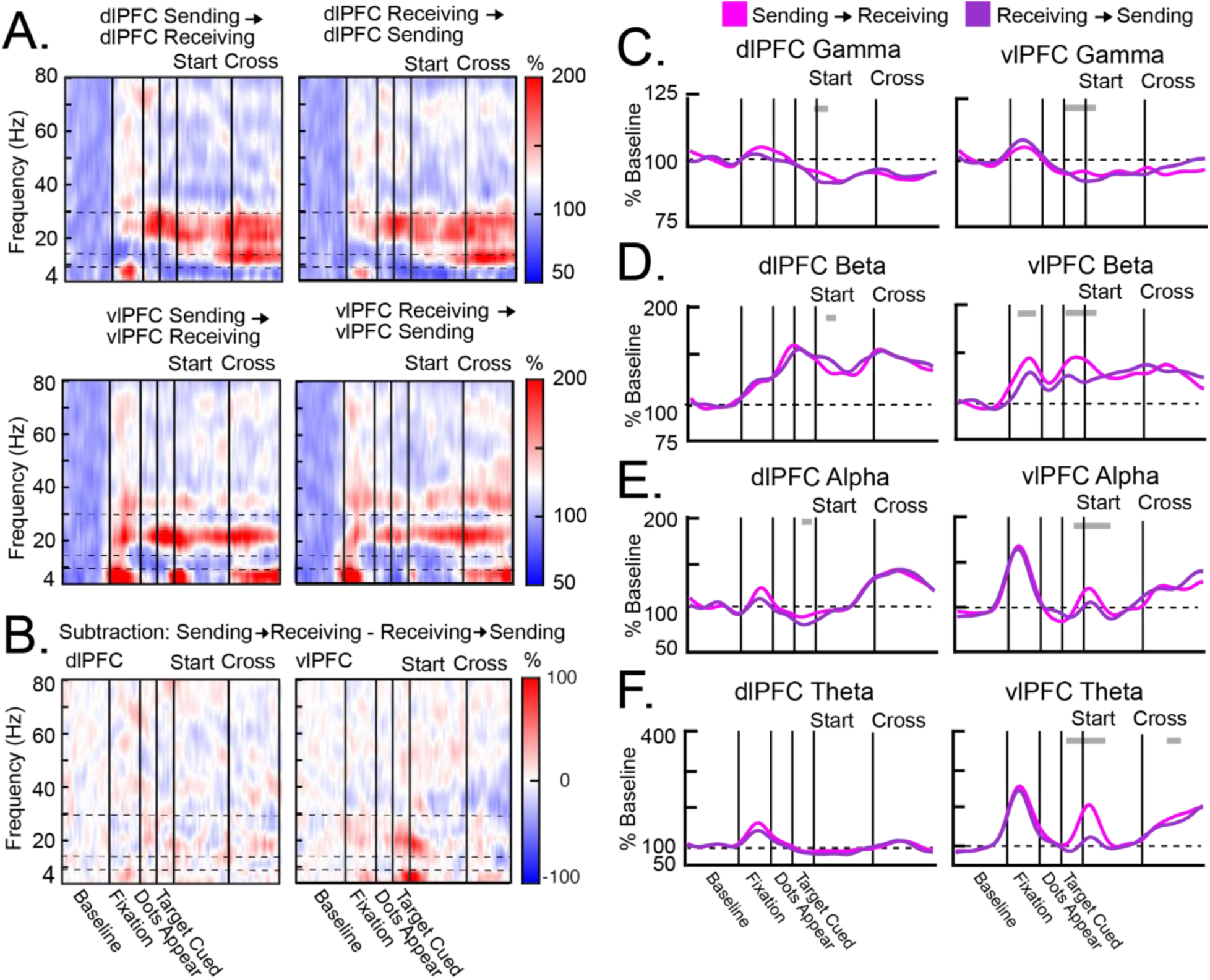
Interhemispheric Granger causality during Hemispheric (Hem) Cross trials. **A,** Heatmaps of interhemispheric Granger causality in the dlPFC (top) and the vlPFC (bottom), from the Sending hemisphere to the Receiving hemisphere (left) and vice versa (right). Horizontal dotted lines indicate the different frequency bands. **B,** Heatmaps of the contrasted Granger causality between hemispheres (i.e., the subtraction of Sending → Receiving causality and Receiving → Sending causality) in the dlPFC (left) and the vlPFC (right). **C-F,** Average Granger causality in the gamma (**C**), beta (**D**), alpha (**E**), and theta (**F**) frequency bands. Granger causality was calculated in the dlPFC (left column) and the vlPFC (right column) separately from the Sending hemisphere to the Receiving hemisphere (pink) and vice versa (purple). All error bars indicate *S.E.M.* Gray bars above each plot indicate time points in which causality was significantly larger in one direction than the opposite direction. Granger causality was largely bidirectional between the Sending and Receiving hemispheres. The exception to this was when the target was cued. In the vlPFC, Sending → Receiving influences were stronger than the opposite direction in all frequency bands, especially for theta and beta.

### Different frequencies had different directions of influence within each hemisphere

Above, we showed that the vlPFC showed stronger bottom-up sensory responses in gamma while the dlPFC showed stronger top-down activity related to the target transfer in alpha/theta frequencies (Fig. 2). This dissociation was also reflected in their direction of influence within each hemisphere. Figures 4A & B show heatmaps of Granger causality between the dlPFC and the vlPFC within each hemisphere (Fig. 4A), as well as the contrast between these directionalities (Fig. 4B). For the first half of the trial, gamma influences were largely bidirectional (Fig. 4C). However, later in the trial, starting before the target cross and continuing after, gamma influences were stronger from the vlPFC to the dlPFC in both the Sending and Receiving hemispheres (Figs. 4B & C; *p* < .001). By contrast, alpha/beta influences were largely in the opposite direction. During target tracking, alpha and beta influences were stronger from the dlPFC to the vlPFC in both hemispheres (Figs. 4B, D & E; *p* < .01). This was also associated with an increase in within-hemispheric alpha/beta coherence during the tracking period (Supplemental Fig. 5). Finally, theta influences were stronger from the vlPFC to the dlPFC than the opposite direction during the initial fixation at the start of the trial (Figs. 4B & F; *p* < .001). This likely reflected the saccade to central fixation. These results suggest that, during tracking, gamma influences tended to flow from the vlPFC to the dlPFC, while alpha/beta influences tended to flow in the opposite direction. This is consistent with the idea that the vlPFC is more associated with bottom-up sensory processing, while the dlPFC is more associated with top-down processing.

**Figure 4.**
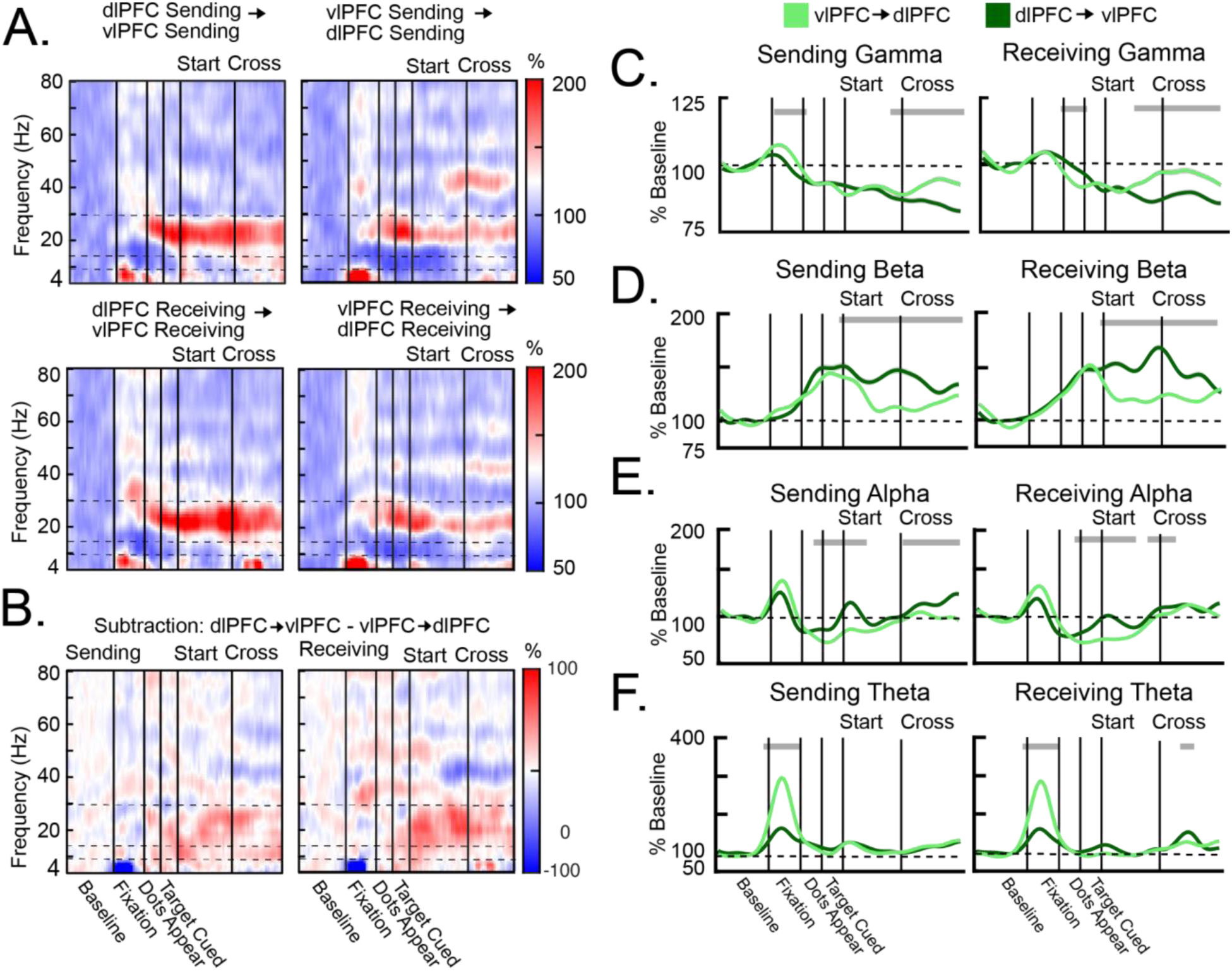
Within-hemispheric Granger causality during Hemispheric (Hem) Cross trials. **A,** Heatmaps of within-hemispheric Granger causality in the Sending hemisphere (top) and the Receiving hemisphere (bottom), from the dlPFC to the vlPFC (left) and from the vlPFC to the dlPFC (right). Horizontal dotted lines indicate the different frequency bands. **B,** Heatmaps of the contrasted Granger causality between prefrontal regions (i.e., the subtraction of dlPFC → vlPFC causality and vlPFC → dlPFC causality). **C–F,** Average Granger causality in the gamma (**C**), beta (**D**), alpha (**E**), and theta (**F**) frequency bands. Granger causality was calculated in the Sending hemisphere (left) and the Receiving hemisphere (right) separately, from the dlPFC to the vlPFC (dark green) and vice versa (light green). All error bars indicate *S.E.M.* Gray bars above each plot indicate time points in which causality was significantly larger in one direction than the opposite direction. During the tracking periods, gamma causality flowed from the vlPFC to the dlPFC, whereas alpha and beta causality flowed from the dlPFC to the vlPFC. Theta causality flowed from the vlPFC to the dlPFC during initial fixation.

### Information about the target was anticipated in the Receiving hemisphere

Spiking activity in both the Sending and Receiving hemispheres carried information about the location of the target. Support Vector Machine (SVM) classifiers were trained to predict the location of the target using spiking activity from each region (Fig. 5A). Classifiers were independently trained and tested at each pair of time points, resulting in a cross-temporal decoding matrix for each region (Fig. 5A). Points along the diagonal reflect “real-time” decoding of target’s location, where the classifiers were trained and tested using the same time point. Points off the diagonal reflect cross-temporal decoding, where classifiers trained at one time point were assayed for their generalization to other time points within the trial. Real-time decoding showed an increase in target location information when the target was cued (Fig. 5B). This information was stronger in the vlPFC than in the dlPFC (*p* < .001), consistent with a stronger role for the vlPFC in bottom-up sensory processing. In the vlPFC, the Sending hemisphere immediately began to carry information about the target’s location as soon as it was cued. By contrast, target information in the Receiving hemisphere ramped up as it approached the hemifield cross. During the transfer, spiking information about the target’s location was high in both hemispheres, suggesting that both hemispheres were tracking the target.

**Figure 5.**
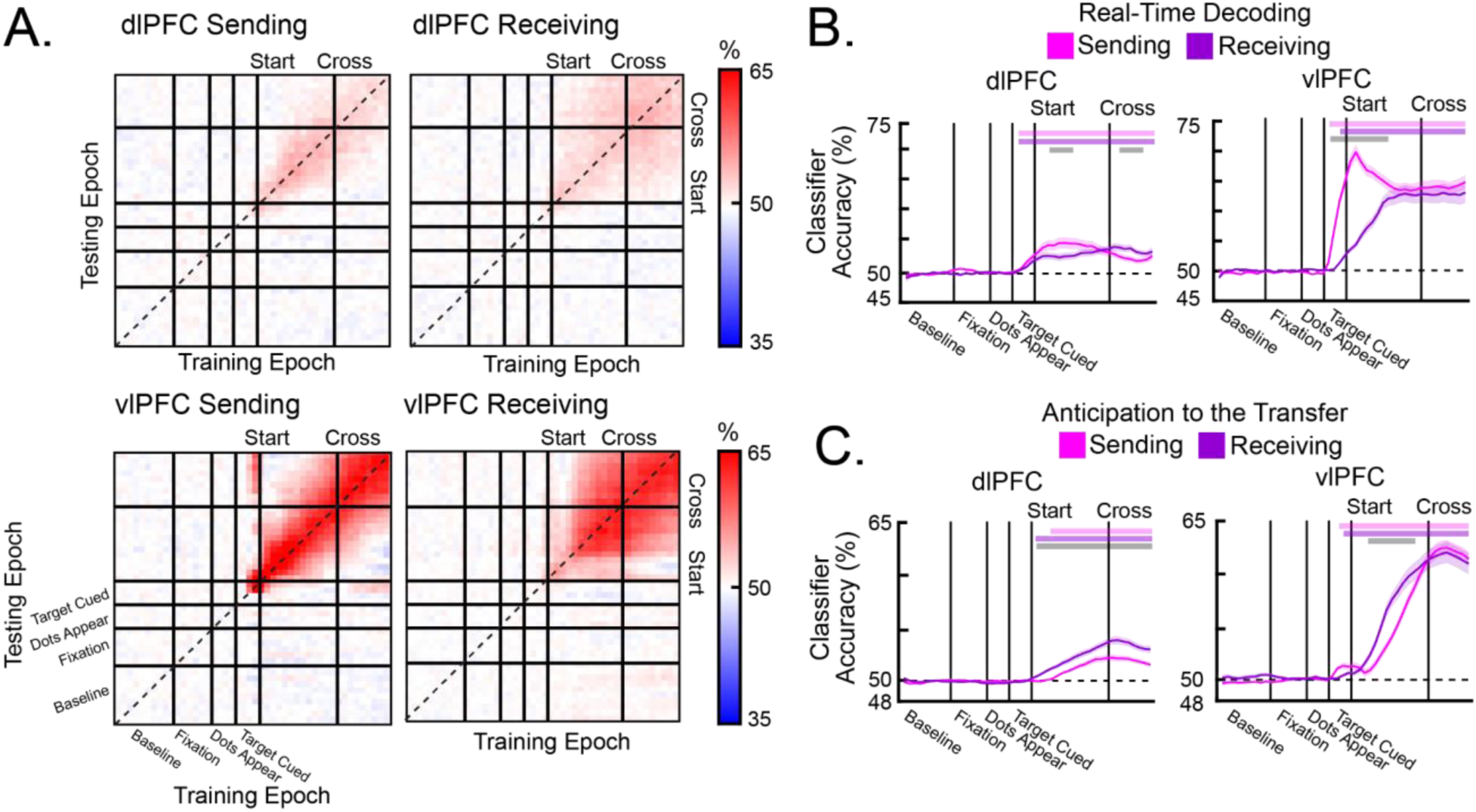
Cross temporal decoding of spiking activity during Hemispheric (Hem) Cross trials. **A,** SVM classifiers were trained to predict the target’s location. Averaged cross-temporal decoding in the dlPFC (top) and the vlPFC (bottom), separated by the Sending hemisphere (left) and the Receiving hemisphere (right). Points on the diagonal reflect real-time decoding. Points off the diagonal reflect cross-temporal generalization. **B,** Classifier accuracies during real-time decoding from classifiers trained with dlPFC activity (left) and vlPFC activity (right). In the vlPFC, classifier accuracy in the Sending hemisphere increased when the target was cued and remained above chance-level throughout the tracking period. Classifier accuracy in the Receiving hemisphere ramped up before the transfer. **C,** Backward-direction cross-temporal generalization from time points after the hemispheric cross, separated by the dlPFC (left) and the vlPFC (right). For both hemispheres, cross-temporal generalization ramped up in anticipation of the cross. All error bars indicate *S.E.M.* Pink bars above each plot indicate time points in which information in the Sending hemisphere was significantly above chance. Purple bars above each plot indicate time points in which information in the Receiving hemisphere was significantly above chance. Gray bars above each plot indicate time points in which classifier accuracies were significantly different between the Sending and Receiving hemispheres.

Cross-temporal decoding revealed that both hemispheres anticipated the cross. Figure 5C shows decoder accuracy when the decoder was trained on neural activity after the cross, then applied backwards in time before the cross. This tests whether the neural code in each hemisphere, after it received or sent the target, generalized to earlier time points in anticipation of the cross. In both the dlPFC and the vlPFC, the Sending and Receiving hemispheres showed significant cross-temporal generalization to time points before the cross (Fig. 5C; all *p* < .01). Thus, the Sending hemisphere seemed to hold onto the target after the target transfer, maintaining it in a similar representation to that prior to target crossing. The Receiving hemisphere instead seemed to activate its post-transfer representation of the target even before it had crossed over into its corresponding hemifield.

## DISCUSSION

Our results provide evidence for an active handoff of attention between cerebral hemispheres. As targets moved between hemifields, their neural representations switched between hemispheres, but not immediately. The Sending hemisphere seemed to hold onto the target after it had crossed hemifields and was thus already sent. The Receiving hemisphere anticipated the target even before it crossed between hemifields. There was a division between bottom-up and top-down processing both anatomically and in neural activity. The vlPFC and higher frequencies (gamma/beta) were more associated with processing bottom-up events such as target cueing and movement. By contrast, the dlPFC and lower frequencies (alpha/theta) were more associated with top-down transfer dynamics that anticipated and signaled the hemispheric transfer. All events, as well as anticipation of the target transfer, were reflected in spiking information. The bottom-up gamma dynamics seemed to flow dorsally from the vlPFC to dlPFC, while top-down alpha/beta flowed ventrally from the dlPFC to the dlPFC. Figure 6 shows a summary of these dynamics.

**Figure 6.**
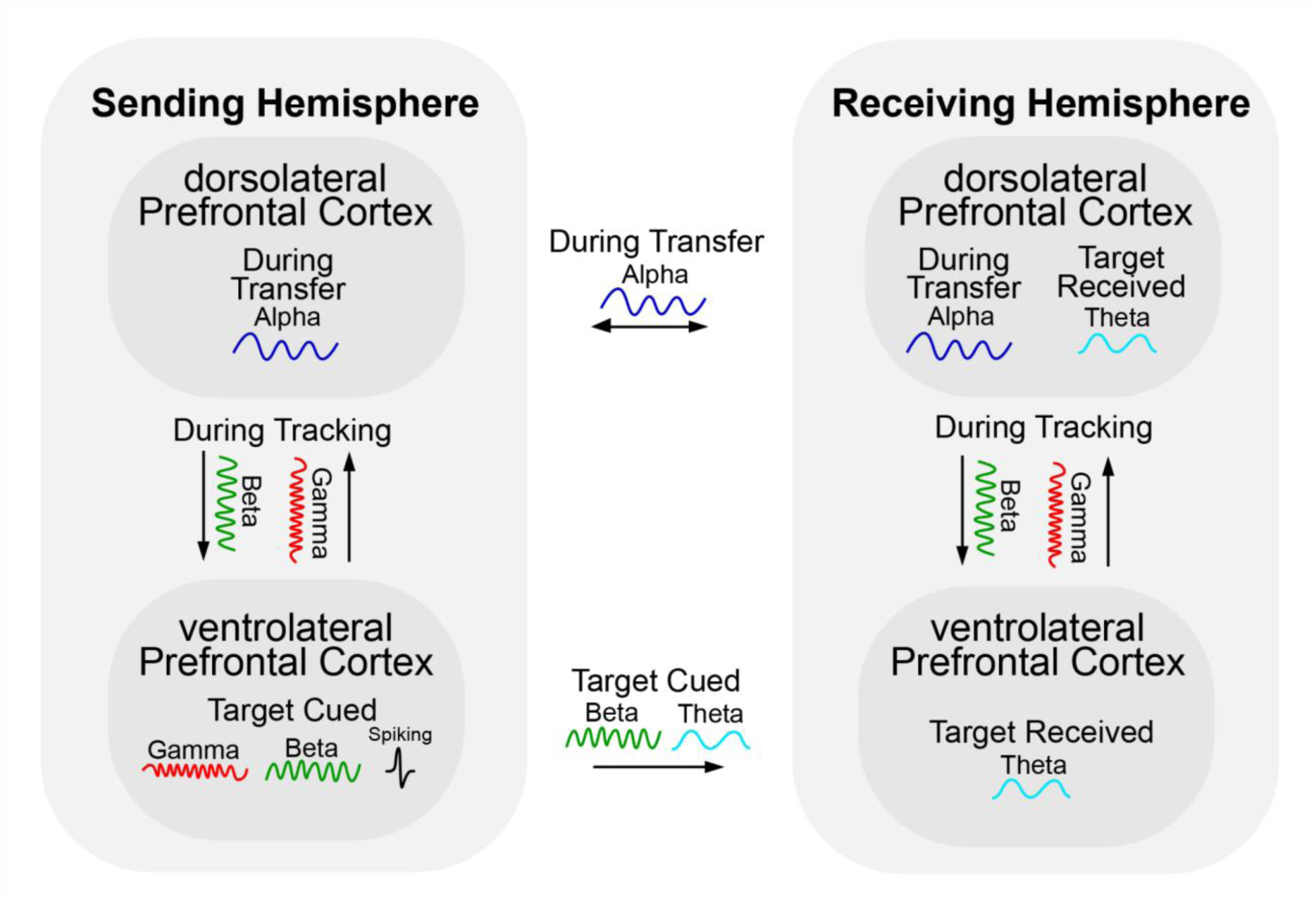
Diagram summarizing the within- and between-hemisphere main effects during Hemispheric (Hem) Cross trials.

Alpha oscillations in cortex have been implicated in spatial attention (Foster et al., 2017; Thut et al., 2006; Worden et al., 2000; van Diepen et al., 2016; Foster et al., 2016; Sauseng et al. 2005; Klatt et al., 2020; Bagherzadeh et al., 2020; Deng et al., 2019; Foster et al., 2017; Sutterer et al., 2021; Wang et al., 2021; Woodman et al., 2021). When attention is directed to a hemifield, there is an increase in alpha in the ipsilateral hemisphere, not the contralateral hemisphere (to which attention is directed). This is consistent with an inhibitory role for alpha in top-down control (Klimesch et al., 2007; Foxe et al., 2011; Klimesch, 2012; Miller et al., 2018; Buschman et al., 2007; Buschman et al., 2012; Jensen et al., 2010). The allocation of alpha can be made in anticipation of a predictable switch in attention or the appearance of a target (Rohenkohl et al., 2011; Samaha et al., 2015; Gould et al., 2011; van Ede et al., 2018; Voytek et al., 2017; Yang et al., 2024). Our alpha effects in the Sending hemisphere could thus be explained by an inhibitory attentional mechanism. As the Sending hemisphere “sends” the target, it is also receiving the distractor that needs to be ignored. However, the alpha effects in the Receiving hemisphere, which is receiving the target, is less straightforward, at least from a spatial attention perspective.

Theta seems to organize networks to reorient attention to new locations/hemifields, resulting in theta power increases at the corresponding location/hemisphere (Helfrich et al, 2019; Fiebelkorn et al., 2018; Fiebelkorn et al., 2019; Voloh et al., 2015; Spooner et al., 2019; Helfrich et al., 2019). This is also consistent with theta power increasing in a hemisphere after working memories are transferred to it (Brincat et al., 2021). We found that theta power increased in the Receiving hemisphere after target transfer. This could reflect attention to the target it now contains.

Beta and gamma oscillations tracked the early sensory events. Beta and gamma were anticorrelated at a fine time scale. Beta decreased as gamma increased. This anticorrelation has been observed in working memory tasks (Lundquist et al., 2016; Lundquist et al., 2018; Lundquist et al., 2023) and occurs when working memories are transferred between hemispheres (Brincat et al., 2021). Beta is thought to inhibit gamma, thus acting as a gating signal for top-down control (Bastos et al., 2015; Jensen et al., 2015; Mendoza-Halloday & Major et al., 2024; Buschman et al., 2007; Richter et al., 2017l van Kerkoerle et al., 2014). The fine-scale push-pull anticorrelation we observed between beta and gamma is consistent with this. We also found that beta/gamma dynamics were stronger in the vlPFC than the dlPFC. This is consistent with evidence from this and other studies that the vlPFC versus the dlPFC are more involved in bottom-up versus top-down processing (Wutz et al., 2018).

These results support an active hand-off of information between hemispheres (Drew et al., 2014). This mechanism is analogous to how modern cell phone towers overlap their spatial coverage (Hegde et al., 2002). As you move from one tower to the next, your call is picked up by the “receiving” tower before it is released by the “sending” tower. This requires more processing but dramatically reduces the number of dropped calls. The brain may utilize a similar handshaking mechanism for interhemispheric transfer (Drew et al., 2014).

Together, these results suggest there are active mechanisms that transfer information between cerebral hemispheres. When visual targets move between the right and left hemifields, they do not merely appear in one hemisphere or the other by virtue of appearance in a hemifield. Instead, the brain seems to anticipate the transfer and acknowledge its completion. Dysfunctional interhemispheric transfer has been associated with schizophrenia (Green, 1978). Many other neurological conditions exhibit reduced coordination between hemispheres, including autism (Anderson et al., 2010), depression (Wang et al., 2015), dyslexia (Dhar et al., 2010), and multiple sclerosis (Cover et al., 2006). Characterizing these hemispheric interactions is important for understanding brain function and disorder.

## Data availability

All data will be made available upon reasonable request.

## Author Contributions

MBB: Conceptualization, Methodology, Investigation, Formal Analysis, Visualization, Validation, Writing-original draft, Writing-review & editing JER: Conceptualization, Methodology, Investigation, Validation, Writing- original draft, Writing- review & editing SLB: Conceptualization, Methodology, Data curation, Software, Methodology, Validation, Writing- original draft, Writing- review & editing MKM: Investigation, Methodology, Writing- review & editing EKM: Conceptualization, Methodology, Funding acquisition, Supervision, Project administration, Writing- original draft, Writing- review & editing

## SUPPLEMENTAL FIGURES

**Supplemental Figure 1.**
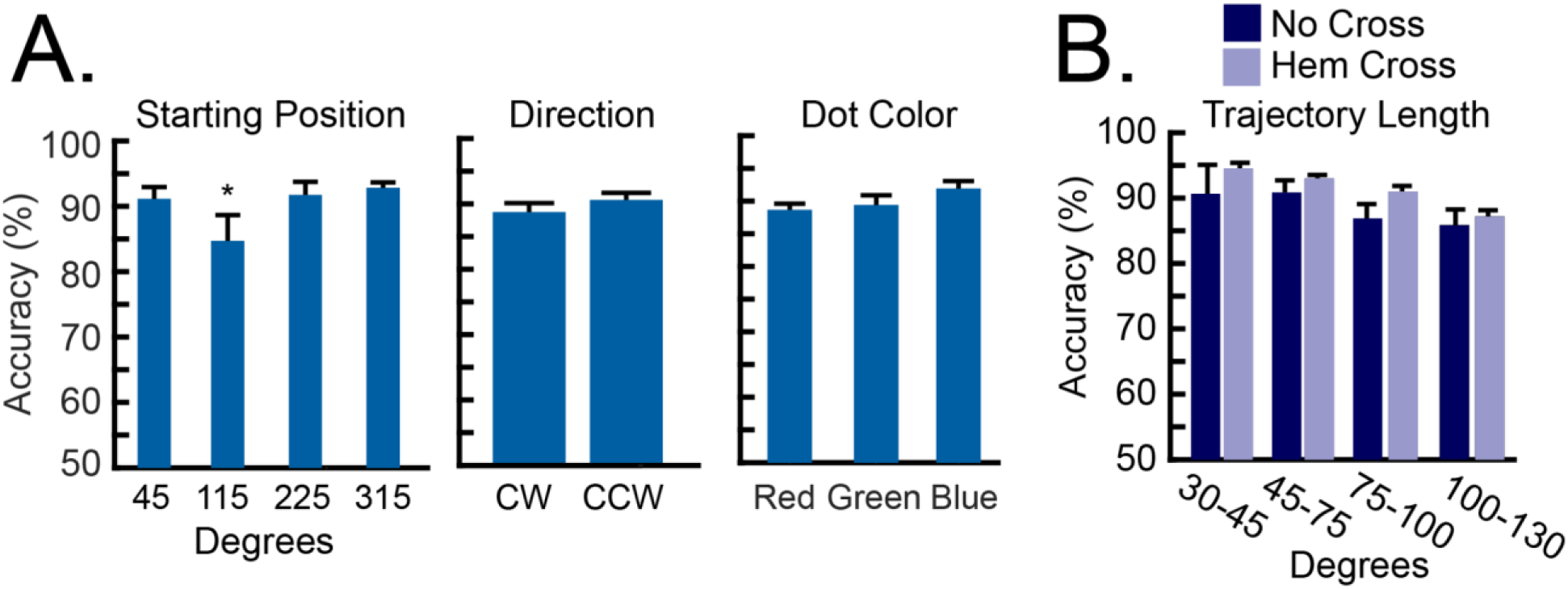
Additional behavioral measures. **A,** Left: Accuracy for each of the four starting positions of the target. The main effect for starting position was significant (*F*(1,28) = 8.28, *p* = .002). Accuracy was significantly lower for the 115-degree position than the other starting positions (*p* < .001). All other comparisons were not significant (*p* > .05). Middle: Accuracy was not significantly different between clockwise (CW) rotation and counter-clockwise (CCW) rotation (*F*(1,28) = 1.90, *p* = .068). Right: Accuracy was not significantly different between the three target colors (i.e., red, green, and blue; *F*(1,28) = 1.57, *p* = .221). **B,** Accuracy for the Hemispheric (Hem) Cross trials and the No control Cross trials, binned at increasing trajectory lengths. There were no significant differences between groups (*F*(2,27) = 2.96, *p* = .069). All error bars indicate *S.E.M*.

**Supplemental Figure 2.**
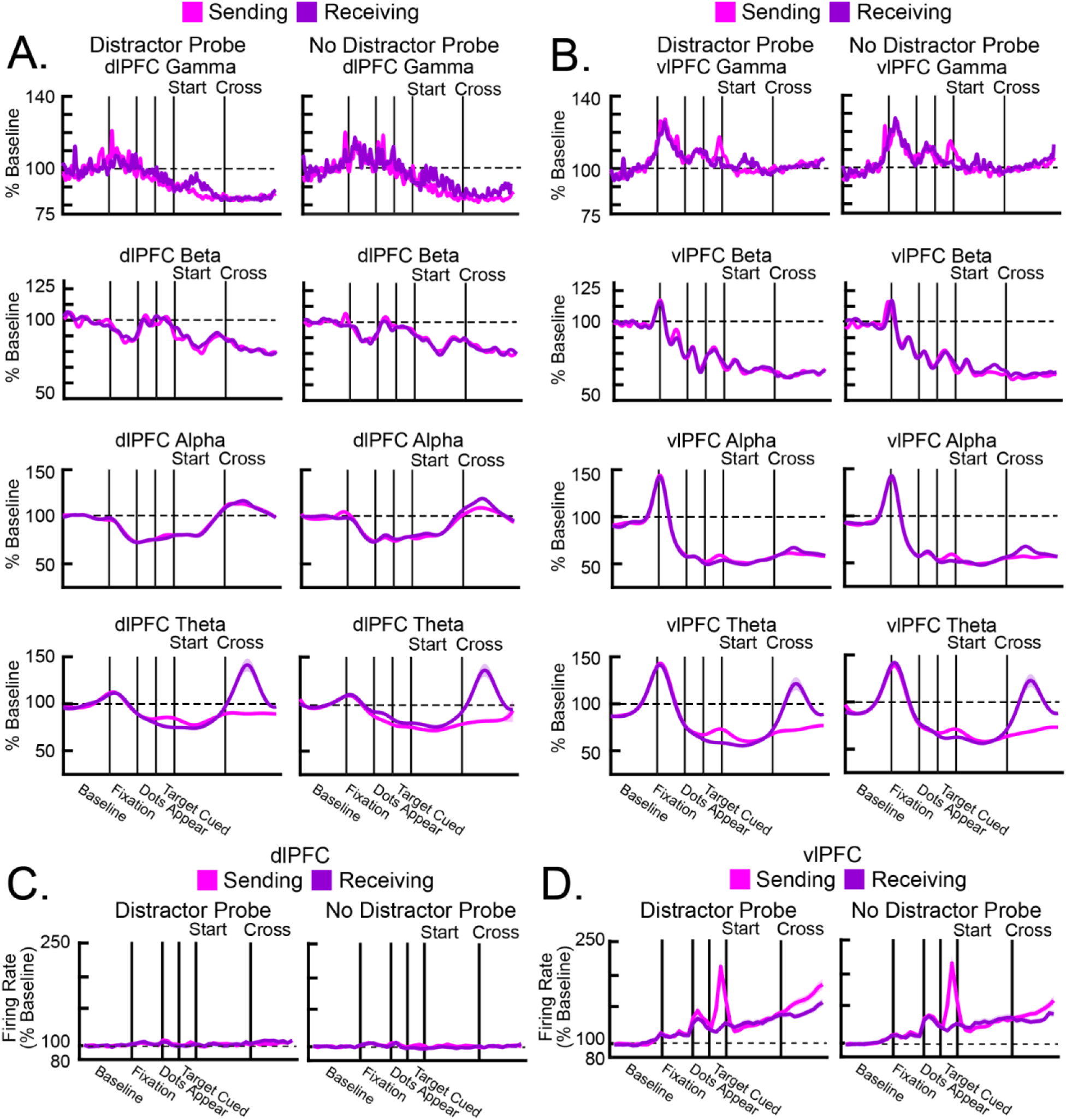
Neural activity was equivalent for trials with and without a color change of the distractor. **A-B,** Average power in the dlPFC (**A**) and the vlPFC (**B**), separated by trials in which the distractor changed color during its trajectory (i.e., Distractor Probe; left) or did not change color during its trajectory (i.e., No Distractor Probe; right). Power was calculated for the Sending hemisphere (pink) and the Receiving hemisphere (purple) separately. In all frequency bands, there were no differences between trial types. **C-D,** Average firing rates in the dlPFC (**C**) and the vlPFC (**D**), separated by the Distractor Probe (left) and No Distractor Probe (right) trials. Firing rates were calculated for the Sending hemisphere (pink) and the Receiving hemisphere (purple), separately. There were no differences in firing rate between trial types. All error bars indicate *S.E.M*.

**Supplemental Figure 3.**
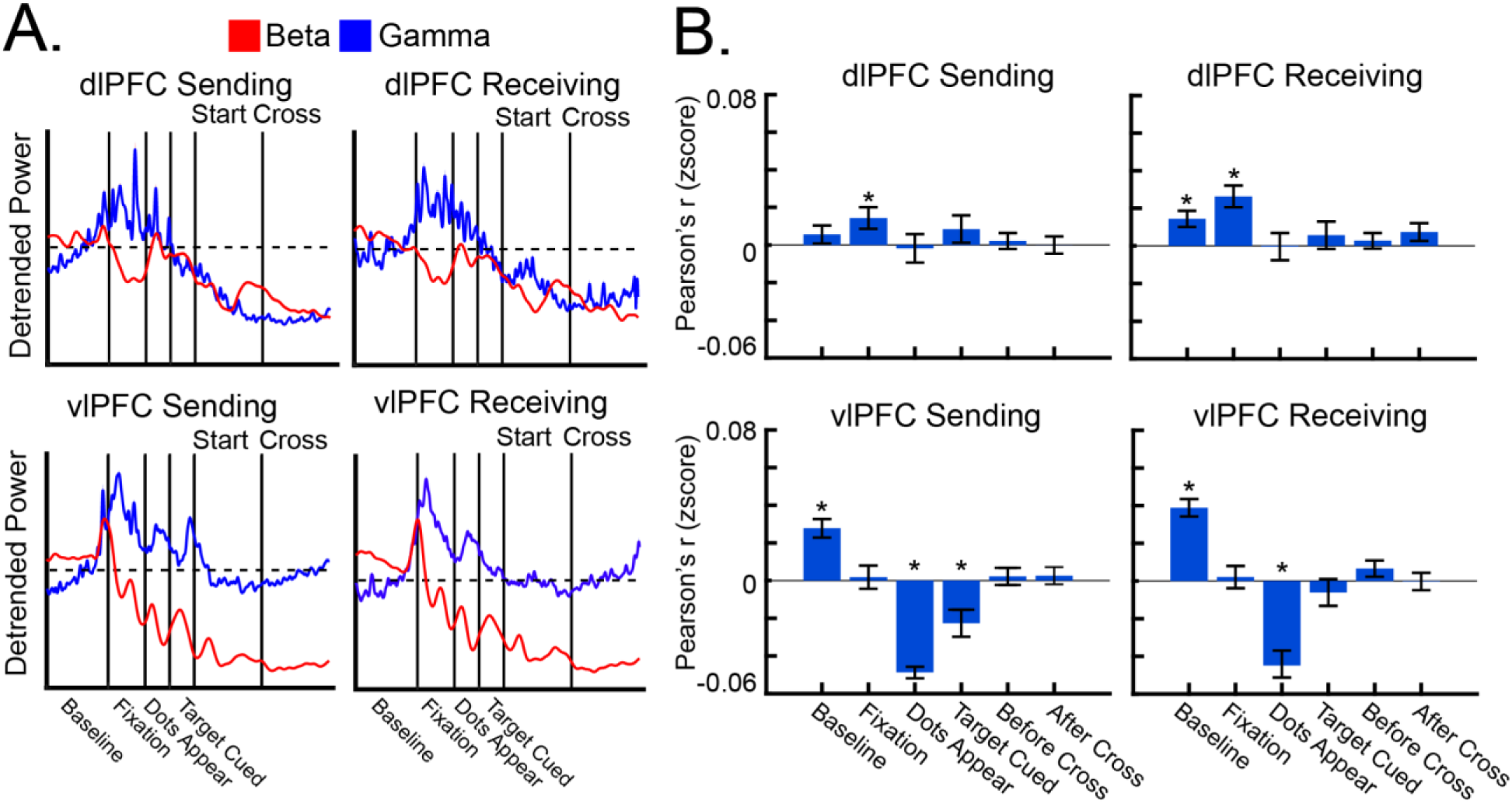
Correlation between beta and gamma power. **A,** Average beta (red) and gamma (blue) power in the dlPFC (top) and the vlPFC (bottom), separated by the Sending hemisphere (left) and the Receiving hemisphere (right). **B,** Average correlation coefficient (with Fisher’s r-to-z transform) between beta and gamma power at each trial event. Correlations were calculated in the dlPFC (top) and the vlPFC (bottom), separated by the Sending hemisphere (left) and the Receiving hemisphere (right). * indicates statistical significance (*p* < .05) against shuffled data. All error bars indicate *S.E.M*.

**Supplemental Figure 4.**
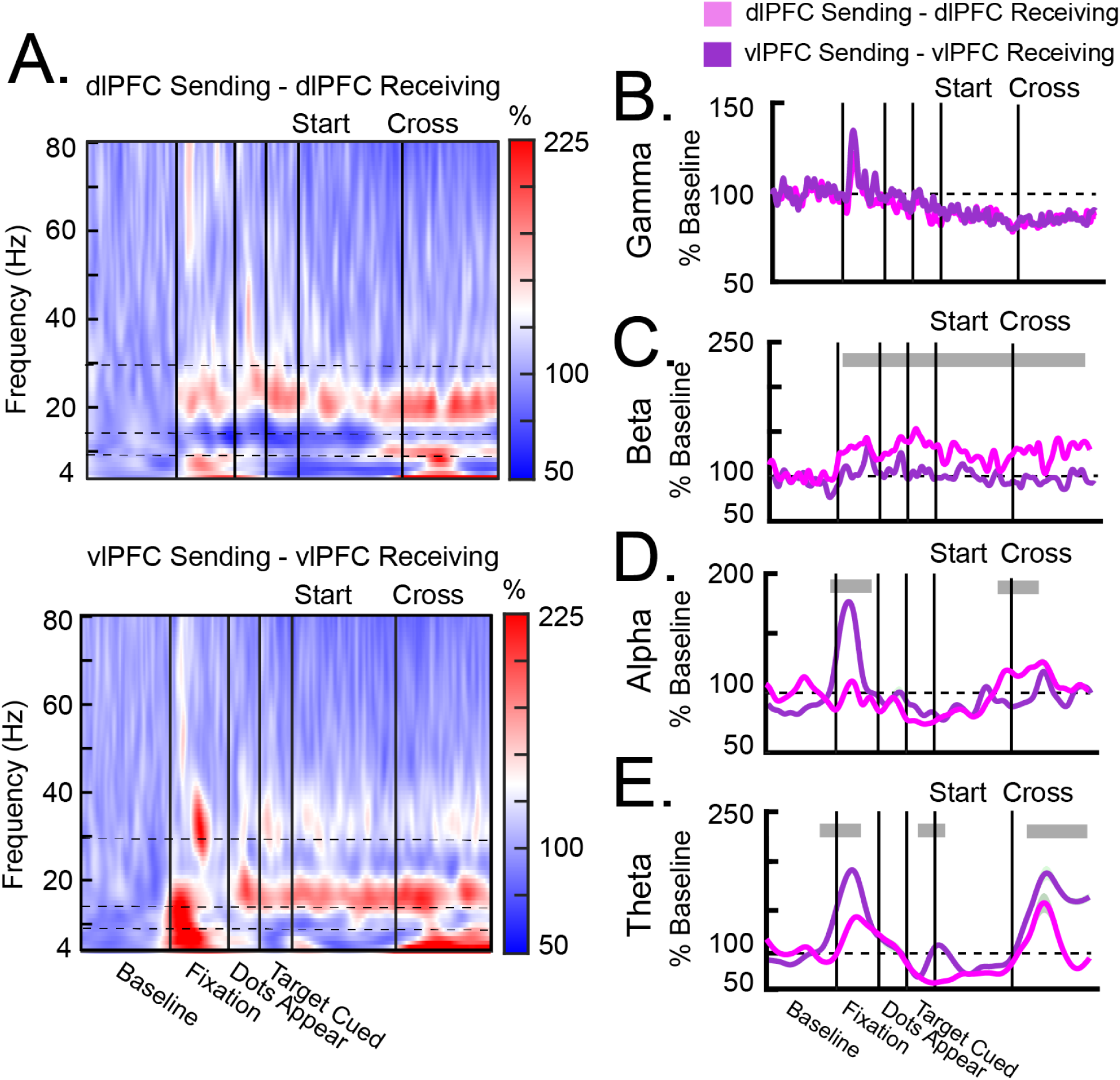
Interhemispheric coherence during Hemispheric (Hem) Cross trials. **A,** Heatmaps of average interhemispheric coherence in the dlPFC (top) and the vlPFC (bottom). Horizontal dotted lines indicate the frequency bands. **B–E,** Average interhemispheric coherence in the gamma (**B**), beta (**C**), alpha (**D**), and theta (**E**) frequency bands. Interhemispheric coherence was calculated in the dlPFC (pink) and the vlPFC (purple) separately. Gray bars above each plot indicate time points in which interhemispheric coherence was significantly different between the dlPFC and the vlPFC. All error bars indicate *S.E.M*.

**Supplemental Figure 5.**
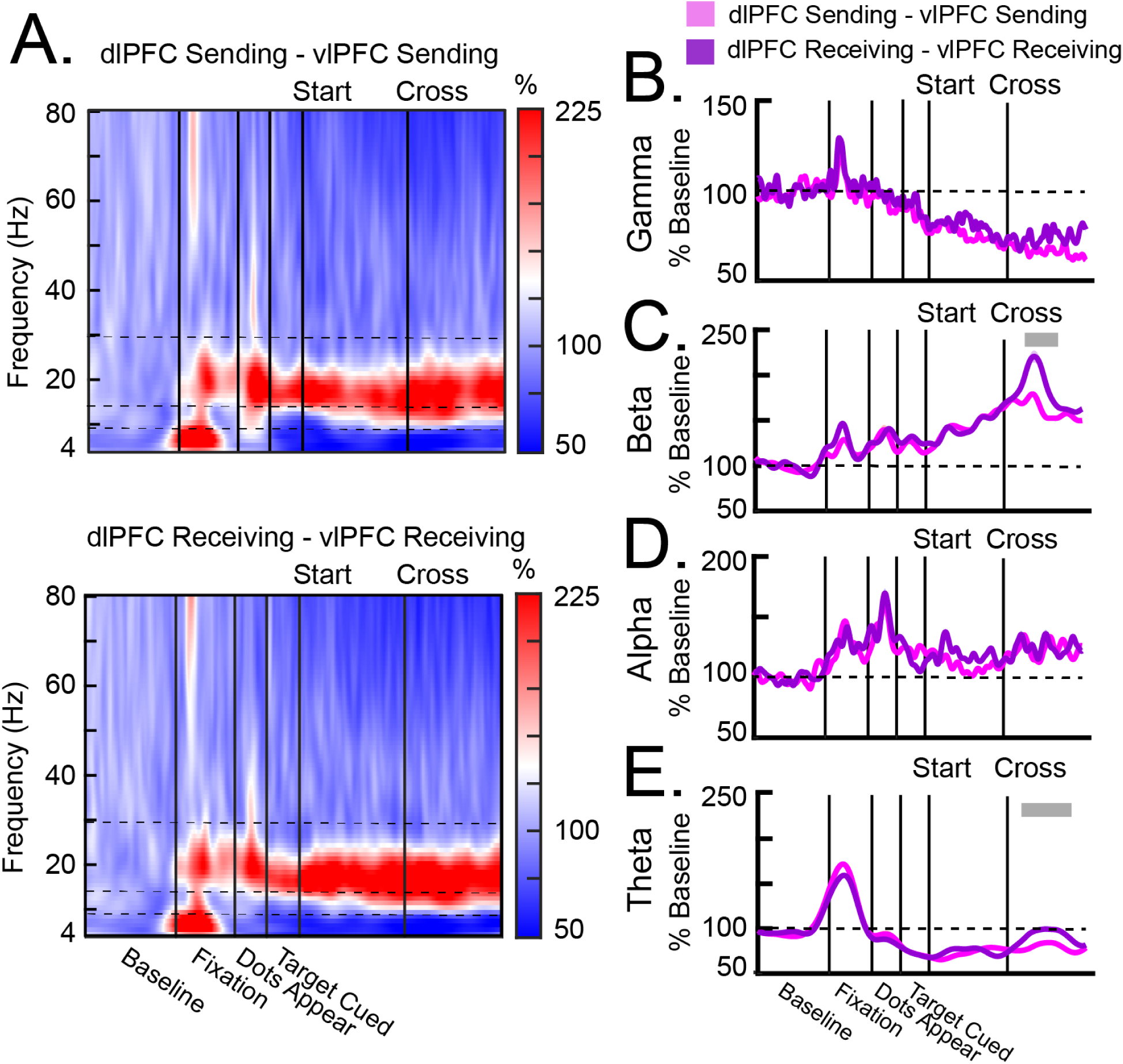
Within-hemispheric coherence during Hemispheric (Hem) Cross trials. **A,** Heatmaps of average within-hemispheric coherence in the Sending hemisphere (top) and the Receiving hemisphere (bottom). Horizontal dotted lines indicate the frequency bands. **B–E,** Average within-hemispheric coherence in the gamma (**B**), beta (**C**), alpha (**D**), and theta (**E**) frequency bands. Within-hemispheric coherence was calculated in the Sending hemisphere (pink) and the Receiving hemisphere (purple) separately. Gray bars above each plot indicate time points in which interhemispheric coherence was significantly different between the dlPFC and the vlPFC. All error bars indicate *S.E.M*.

